# *Arc-*Expressing Accessory Olfactory Bulb Interneurons Support Chemosensory Social Behavioral Plasticity

**DOI:** 10.1101/2022.05.02.490296

**Authors:** Kelsey E. Zuk, Hillary L. Cansler, Jinxin Wang, Julian P. Meeks

## Abstract

The accessory olfactory system (AOS) is critical for the development and expression of social behavior. The first dedicated circuit in the AOS, the accessory olfactory bulb (AOB), exhibits cellular and network plasticity in male and female mice after social experience. In the AOB, interneurons called internal granule cells (IGCs) express the plasticity-associated immediate-early gene *Arc* following intermale aggression or mating. Here, we sought to better understand how *Arc*-expressing IGCs shape AOB information processing and social behavior in the context of territorial aggression. We used “ArcTRAP” (Arc-CreERT2) transgenic mice to selectively and permanently label *Arc*-expressing IGCs following male-male resident-intruder interactions. Using whole-cell patch clamp electrophysiology, we found that *Arc*-expressing IGCs display increased intrinsic excitability for several days after a single resident-intruder interaction. Further, we found that Arc-expressing IGCs maintain this increased excitability across repeated resident-intruder interactions, during which resident mice increase or “ramp” their aggression. We tested the hypothesis that *Arc*-expressing IGCs participate in ramping aggression. Using a combination of ArcTRAP mice and chemogenetics (Cre-dependent hM4D(G_i_)-mCherry AAV injections), we found that disruption of Arc-expressing IGC activity during repeated resident-intruder interactions abolishes the ramping aggression exhibited by resident male mice. This work shows that *Arc*-expressing AOB IGC ensembles are activated by specific chemosensory environments, and play an integral role in the establishment and expression of sex-typical social behavior. These studies identify a population of plastic interneurons in an early chemosensory circuit that display physiological features consistent with simple memory formation, increasing our understanding of central chemosensory processing and mammalian social behavior.

**Significance statement:** The accessory olfactory system (AOS) plays a vital role in rodent chemosensory social behavior. We studied experience-dependent plasticity in the accessory olfactory bulb (AOB) and found that internal granule cells (IGCs) expressing the immediate-early gene *Arc* after the resident-intruder paradigm increase their excitability for several days. We investigated the roles of these Arc-expressing IGCs on chemosensory social behavior by chemogenetically manipulating their excitability during repeated social interactions. We found that inhibiting these cells eliminated intermale aggressive ramping behavior. These studies identify a population of plastic interneurons in an early chemosensory circuit that display physiological features consistent with simple memory formation, increasing our understanding of central chemosensory processing and mammalian social behavior.

## Introduction

The survival and reproduction of an animal is highly dependent on its ability to appropriately interact with its social environment. Social interactions arise from the integration of complex sensory cues found in the animal’s surroundings. More so than humans, terrestrial mammals rely on the detection and interpretation of olfactory stimuli in order to guide their behavior, both during initial encounters and after experience (Brennan, et al., 2015). In rodents, the accessory olfactory system (AOS) plays an essential role in guiding behavior through the detection of chemosignals, which are thought to convey information about the species, sex, reproductive status, or even health of other animals (Brennan, 2001; Mucignat-Caretta, et al., 2012; Salazar, et al., 2001). Many chemosignals, including pheromones, are detected in the AOS via vomeronasal sensory neurons (VSNs) found in the vomeronasal organ (VNO). VSNs are known to be activated by excreted proteins, peptides, and steroids (Chamero, et al., 2007; Kimoto, et al., 2005; Leinders-Zufall, et al., 2004; Liberles, et al., 2009; Nodari, et al., 2008; Riviere, et al., 2009; Wong, et al., 2020). Disruption of VSN signaling through knockout of the TRPC2 ion channel has been shown to alter the display of sex-typical behavior, such as mating and aggression, reinforcing the notion that sensory input to the AOS plays a critical role in behavioral output (Stowers, et al., 2002). Overall, it is well-established that AOS alteration disrupts social behavior, but the circuit-level mechanisms mediating these behavioral changes are still not understood.

Previous work suggested that inhibitory signaling, via GABAergic neurons, plays an essential role in regulating olfactory information processing (Brennan, et al., 1997; Jia, et al., 1999). Recent studies identified a population of GABAergic inhibitory interneurons in the accessory olfactory bulb (AOB), the first dedicated circuit for AOS information processing, that display increased intrinsic excitability for several hours after social chemosensory experience (Cansler, et al., 2017; Gao, et al., 2017). These inhibitory interneurons, known as internal granule cells (IGCs), are found in the internal cellular layer (ICL) of the AOB and are the most abundant inhibitory interneuron subtype in this circuit (Larriva-Sahd, 2008). IGCs have small basal dendrites that reside in the ICL and long apical dendrites that stretch into the external cellular layer (ECL), where excitatory mitral cells reside. Each IGC forms reciprocal, dendro-dendritic synapses along the dendrites of multiple mitral cells (Larriva-Sahd, 2008; Taniguchi, et al., 2001). These cells are analogous to granule cells in the main olfactory bulb (MOB), which have similar morphology, connectivity, and presumably function to their AOB counterparts (Sailor, et al., 2016). MOB granule cells demonstrate changes in their dendritic arbors and dendro-dendritic synapses throughout the life of an animal (Quast, et al., 2017; Sailor, et al., 2016). Experience-dependent structural changes, as measured by electron microscopy, in these dendro-dendritic synapses have also been shown to be associated with olfactory learning in the AOB (Matsuoka, et al., 1997). Building on recent work, we sought to further understand how AOB experience-dependent plasticity may be linked to changes in immediate-early gene (IEG) expression in inhibitory interneurons (Cansler, et al., 2017; Gao, et al., 2017).

IEGs, which are quickly and selectively upregulated after a neuronal stimulus response, have been linked to various forms of plasticity (Minatohara, et al., 2015). The activity-regulated cytoskeletal-associated gene (*Arc)* has been shown to play a direct role in mediating long-term depression (LTD) in hippocampal excitatory neurons (Bramham, et al., 2008; Jakkamsetti, et al., 2013; Shepherd, et al., 2011). AOB IGCs that display increased excitability after experience also selectively express *Arc* mRNA and protein (Arc) (Cansler, et al., 2017). Arc is associated with experience-dependent plasticity throughout the brain (reviewed in Korb, et al., 2011), and we hypothesized that Arc-expressing IGCs contribute to AOB function in the context of AOS-mediated social behavior.

Here, we report that IGCs that transiently express *Arc* following a single social experience display increased intrinsic excitability for several days. Increased IGC excitability persisted during a week of repeated social experiences, but did not scale with the number of experiences. We tested whether Arc-expressing IGCs participate in the development of aggressive behaviors between males, an AOS-mediated behavior. We found that chemogenetically inhibiting Arc-positive AOB IGCs completely abolished aggressive ramping over multiple intermale resident-intruder interactions. Our results indicate that this early sensory inhibitory neuron population plays a critical role in regulating the behavioral response to social chemosignals.

## Materials and Methods

### Mice

All animal procedures were conducted in compliance with both the University of Texas Southwestern Medical Center Institutional Animal Care and Use Committee (IACUC) and the University of Rochester University Committee on Animal Resources (UCAR). Sexually naïve adult male mice aged 6-12 weeks were used throughout the study. Mice were housed on a reverse 12-hour light/dark cycle, with lights on from either noon to midnight or 6pm-6am. Food and water were provided *ad libitum*. B6.129(Cg)-Arctm1.1 (ArcCreERT2 +/-; the Jackson Laboratory; Stock #021881) were crossed with Gt(ROSA)26Sortm9(CAG-tdTomato)Hze (Ai9 +/- or +/+; the Jackson Laboratory; Stock #007905) (Guenthner, et al., 2013). These mice will be referred to as ArcTRAP (ArcCreERT2 +/-, Ai9 +/- or +/+) throughout (Guenthner, et al., 2013). In some experiments, the ArcTRAP line was crossed with a hemizygous Arc-d4EGFP line (a gift from Dr. Pavel Osten, via Dr. Kimberly Huber) (Grinevich, et al., 2009) to form ArcTRAP x Arc-d4EGFP mice. Each mouse line was maintained on a C57BL/6J background. BALB/cJ mice were obtained from either the University of Texas Southwestern Mouse Breeding Core or the Jackson Laboratory (Stock #000651). A total of 207 mice were used in this study.

### Resident-intruder test

All behavioral tests were conducted during the dark cycle. ArcTRAP resident male mice were single housed for one week without cage changes. At the end of one-week, resident mice underwent a process referred to as “TRAP” (targeted recombination in active populations) (Guenthner, et al., 2013), where they were lightly anesthetized with isoflurane (to reduce stress from injection) and intraperitoneally injected with 4-hydroxytamoxifen (4-OHT; Sigma-Aldrich) at a dose of 50 mg/kg. Twenty minutes after 4-OHT injection, a BALB/cJ intruder male was introduced to the residents’ cage for 10 minutes during the resident-intruder test. After 10 minutes, the intruder was returned to its home cage. Resident mice were left in their home cage and allowed to rest for at least 1 hour before being returned to the animal housing facility.

The resident-intruder test was conducted once per day, for up to 8 days. All behavior sessions were recorded with handheld cameras (Panasonic). Aggressive behaviors were quantified by manual video review and DeepLabCut (Mathis, et al., 2018). DeepLabCut is a computer software package used for animal pose estimation and is based on transfer learning with deep neural networks (Mathis, et al., 2020). For a subset of our DREADD-manipulated and control mice, behavior videos were recorded from both a side view (used for all manual behavior annotation) and a top view (better suited for DeepLabCut). DeepLabCut was trained with 1184 video frames that were manually annotated, and 12 body labels were applied to the resident and intruder. From the manual annotation, we identified 20 resident behaviors that could be analyzed for state duration (occupancy). Overall, 259 behavioral features/metrics were calculated based on the tracking data generated from DeepLabCut, including: distance, velocity, body orientation, and position. A random forest classifier was applied to classify the 20 behaviors of interest. After training, the performance of the random forest classifier was evaluated by 5-fold cross-validation, and the accuracy of the final classifier was 0.943. For manual video analysis, the number of resident-initiated aggressive interactions were counted. These aggressive interactions included attack behavior (lunging at the intruder with or without biting) and fighting behavior (active physical altercation between the resident and intruder, often involving tumbling). Sequential behaviors were counted separately if a physical separation occurred between the resident and intruder, or if there was at least a 2 second pause in the interaction.

### Live slice preparation

Mice were anesthetized with isoflurane and decapitated 1, 3, 5, or 7 days after being TRAPed on Day 0. Brains were dissected, and 400 μm parasagittal sections of the AOB were prepared using a vibrating microtome (Leica VT1200) in ice-cold, oxygenated ACSF. The ACSF contained (in mM) 125 NaCl, 2.5 KCl, 2 CaCl_2_, 1 MgCl_2_, 25 NaHCO_3_, 1.25 NaH_2_PO_4_, 25 glucose, 3 myoinositol, 2 Na-pyruvate, and 0.4 Na-ascorbate, with an additional 9 MgCl_2_ in the slicing buffer. After slicing, the slices were kept in a recovery chamber at room temperature (23 °C) containing oxygenated ACSF with 0.5 mM kynurenic acid to prevent potential glutamate excitotoxicity during the recovery/holding period. Just before recordings, slices were transferred to a slice chamber (Warner Instruments) mounted on an upright fluorescence-equipped differential interference contrast microscope (Nikon; Model FN1). The tissue was superfused with oxygenated ACSF at 35°C via a peristaltic pump (Gilson) at a rate of 3-3.5 ml/min.

### Electrophysiology

Whole-cell patch-clamp recordings were made on ArcTRAP+ (tdTomato+) and ArcTRAP- (tdTomato-) IGCs 1, 3, 5, or 7 days following TRAP on Day 0. Thin wall borosilicate glass electrodes with a tip resistance between 4.5 and 13.5 MΩ were filled with internal solution containing the following (in mM): 115 K-gluconate, 20 KCl, 10 HEPES, 2 EGTA, 2 MgATP, 0.3 Na_2_GTP, 10 Na phosphocreatine at pH 7.37. All recordings were amplified using a MultiClamp 700B amplifier (Molecular Devices) at 20 kHz and were digitized by a DigiData 1440 analog-digital converter controlled via pClamp 10.5 software (Molecular Devices; RRID:SCR_011323). Data were analyzed by Clampex 10.5 (Molecular Devices) and custom software written in MATLAB.

Patched AOB IGCs were subjected to a series of current-clamp and voltage-clamp challenges. Immediately after achieving the whole-cell configuration, the resting membrane potential (V_rest_) of each cell was measured in current-clamp mode. To standardize measurements across cells with different V_rest_, we injected steady-state currents to maintain the membrane potential (V_m_) of each cell between −70 and −75 mV. Based on initial measurements of input resistance (R_input_), we empirically determined the amplitude of hyperpolarizing current that adjusted V_m_ by −50 mV (to approximately −125 mV). After determining this initial current injection amplitude, we generated a cell-specific 10-sweep Clampex protocol that applied increasingly depolarizing 0.5 s square current pulses, starting with the initial injection amplitude. For example, if the initial current injection was determined to be −100 pA, the 10-sweep protocol would have current injection increments of +20 pA (i.e., −100, −80, −60, … +80 pA). In voltage clamp, cells were initially held at −70 mV, and a series of 12 voltage command steps (0.5 s in duration) were applied that spanned −100 to +10 mV. For each cell, both current-clamp and voltage-clamp protocols were applied up to four times, and all reported quantities represent the mean responses across repeated trials.

For CNO wash-on experiments, a 32-minute recording was conducted while measuring baseline resting membrane potential. tdTomato signal was used to identify ArcTRAP+ IGCs in slices that contained either the AAV-Gi-DREADD or the AAV-mCherry. In the first two minutes of the recording, ACSF was washed over the slice. In minutes 2-12, 10 µM CNO in ASCF was washed-on. ACSF was then washed over the slice for the remainder of the recording, between minutes 12-32.

### Injectable drug preparation

4-OHT (Sigma; Catalog #H6278) was freshly dissolved in warmed ethanol at a concentration of 20 mg/ml on each experimental day. Once dissolved, a volume of corn oil (Acros Organics; Catalog #8001-30-7) equivalent to the volume of ethanol was added. The mixture was heated and spun at 1725 RPM in a Speedvac concentrator (Savant; Model #SVC-100H) for 20 minutes. The 4-OHT/corn oil solution was then loaded into insulin syringes (BD; Catalog #329461) at a dose of 50mg/kg for injection. Clozapine-n-oxide (CNO; Tocris; Catalog #4936) was dissolved in 0.5% DMSO (Sigma; Catalog #D2438) (v/v in 0.9% sterile saline) at RT to a concentration of 3.79 mg/ml. The solution was then aliquoted and stored at -20°C until experimentation. A dose of 2.5mg/kg was used for injection. 0.5% DMSO (v/v in 0.9% sterile saline) was used as a vehicle control injection.

### Viral injections for DREADD experiments

pAAV-hSyn-DIO-hM4D(Gi)-mCherry (Addgene; Catalog #44362-AAV9; RRID: Addgene_44362) or pAAV-hSyn-DIO-mCherry (Addgene; Catalog #50459-AAV9; RRID: Addgene_50459) was utilized for bilateral intracranial injections into the accessory olfactory bulb (AOB) of resident ArcTRAP mice (Krashes, et al., 2011). Mice were head-fixed on a custom stereotaxic device that allows for a 30-degree tilt along the horizontal plane, elevating the rostral brain. A Nanoject II (Drummond Instruments) was used to deliver a volume of 92-110nL at coordinates AP +3800 µm, ML +/- 1000 µm, DV -2000 µm relative to Bregma. After one week of recovery, mice were TRAPed. The mice were then given another week to allow for expression of the Cre-dependent G_i_-DREADD. Subsequently, the mice were exposed to repeated resident intruder interactions following intraperitoneal injections with either DMSO (control; 0.5% DMSO v/v in 0.9% sterile saline) or clozapine-n-oxide (CNO; 2.5mg/kg in 0.5% DMSO).

### Tissue Imaging

Fixed tissue imaging was conducted at the URMC Histology, Biochemistry, and Molecular Imaging core facility using an Olympus VS120 Virtual Slide Microscope and Visiopharm Image Analysis System. Resident mice were TRAPed on Day 0 where they received either 4-OHT and the resident-intruder assay, 4-OHT alone, or the resident-intruder assay alone. On Day 3, mice were euthanized with ketamine/xylazine (300 mg/kg / 30mg/kg) and intracardially perfused with 0.9% sodium chloride, followed by 4% paraformaldehyde (PFA). Brains were dissected and post-fixed in 4% PFA for 24-48 hours. 30% sucrose was used to cryoprotect the brains overnight before embedding in OCT compound (Fisher) for cryosectioning on a Shandon Cryotome FSE (Thermo Electron Corporation). Sections were cut at 25 µm thickness and free-floated in PBS prior to mounting. Sections were mounted using Vectashield Hardset antifade mounting medium (Vector Laboratories). 20X Z-stack (15 slices, 1.34um spacing) images were captured in the TRITC channel (580nm emission wavelength). Images were first opened in QuPath (Bankhead, et al., 2017) and converted to .tiff format for processing in FIJI/ImageJ (Schindelin, et al., 2012). Maximum intensity projections were used, and brightness/contrast was adjusted equally for all images. A threshold was set to include tdTomato+ cell bodies and the watershed algorithm was applied to separate cell bodies in close proximity. A region of interest (ROI) was drawn around the ICL of the AOB. Particles were included based on size and circularity, and analyzed by the built in “analyze particles” ImageJ function. Cell counts were normalized to the ROI area.

Live tissue 2-photon imaging was conducted on ArcTRAP-d4EGFP mice. Images were captured on a Thorlabs Acerra upright 2-photon microscope system equipped with an XLUMPlanFLN 20X objective (Olympus) and a fast-scanning resonant galvanometer along one of the two principal axes. ArcTRAP-d4EGFP residents were TRAPed on Day 0. On days 1 and 2, they were re-exposed to the intruder seen on Day 0. On Day 3, residents were either re-exposed to the same intruder seen previously, or exposed to a novel female. Three hours after the Day 3 resident-intruder interaction, brains were acutely dissected and live slices were prepared (as described above). 200 µm thick image stacks were taken from the AOB with ∼0.5 x 0.5 x 2 µm sampling, with excitation at 890 nm and emission filters for eGFP (525 ± 50 nm) and tdTomato (607 ± 70 nm). Each image was generated by averaging 16 sequentially acquired frames. Images were stitched using ImageJ (Schindelin, et al., 2012). Maximum intensity projections were used. Histogram-matching bleach correction and background subtraction were applied to each image (Miura, 2020). An ROI was drawn around the ICL of each image and the colocalization threshold function was applied. The Manders coefficients with applied thresholds were calculated for both red and green channels and reported (Manders, et al., 1993; Fig 3C-D).

## Results

### Arc expression is upregulated in AOB IGCs after the resident-intruder paradigm

To study Arc-expressing IGCs over week-long time periods, we first wanted to validate the use of the ArcTRAP transgenic mouse line. This transgenic mouse contains a tamoxifen-inducible Cre directed by the *Arc* promoter. When *Arc* expression occurs, it causes expression of tamoxifen-inducible Cre. If tamoxifen is present, Cre will excise the stop codon on the Ai9 transgene, permanently labeling that neuron with tdTomato. We tested whether tdTomato expression was elevated in mice after inducing Cre with 4-OHT and subjecting male mice to the resident-intruder assay, which has been used to model male-male territorial aggression. Resident male ArcTRAP mice were solo-housed for one week prior to behavioral testing. On Day 0, resident mice received an injection of 4-OHT and were left in their home cage to habituate for 20 minutes. Following the 20-minute habituation period, a male BALB/cJ intruder mouse was introduced to the resident’s home cage for 10 minutes (Fig. 1A). This combination of solo-housing and 4-OHT injection/behavior on Day 0 will be referred to as “TRAP” (targeted recombination in active populations) throughout the text. As expected, residents that received both the 4-OHT injection and the resident-intruder assay showed a significant increase in tdTomato expression in the ICL of the AOB, compared to residents that received either the 4-OHT injection or resident-intruder assay alone (one-way ANOVA, p=0.0059 and p<0.0001, respectively; Fig. 1B-E). Because Arc is induced in AOB IGCs through the detection of accessory olfactory chemosignals in the mouse’s environment (Cansler, et al., 2017), we were not surprised to see some tdTomato expression in those residents only receiving the 4-OHT injection. This indicates that the residents are able to detect their own chemosignals in their home cage, and that this detection is enough to induce some level of Arc expression in select IGCs, or that there is a baseline expression of Arc in IGCs. Overall, these results support the use of the ArcTRAP transgenic mouse line for measuring and tracking Arc-expressing IGCs across time in the context of male-male territorial aggression.

**Figure 1.**
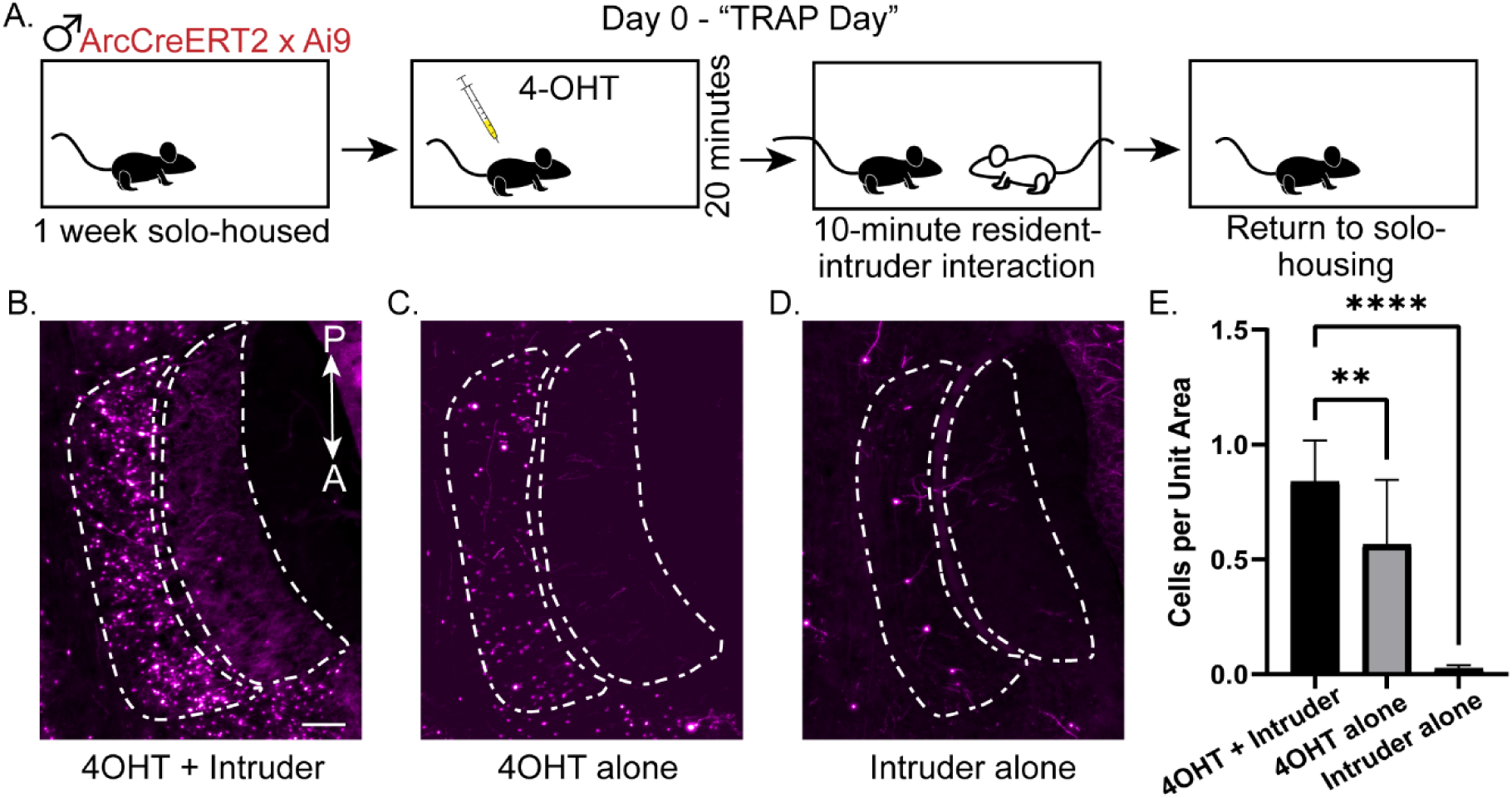
TRAPing during resident-intruder interactions permanently labels AOB IGCs with tdTomato. (**A**) Depiction of the TRAPing process (paired 4-OHT injection and social interaction). Day 0 is “TRAP Day.” (**B-D**) Sample AOB images of tdTomato expression in ArcTRAP mice in “4-OHT+Intruder” (**B**), “4-OHT alone” (**C**), and “Intruder alone” (**D**) conditions. In all three images, the ICL is outlined by the dashed region to the left, and the external cellular layer and glomerular layer are outlined by the dashed region to the right. Scale bar: 100 µm. (**E**) Quantification of tdTomato-positive cells in the ICL. Cell counts are normalized to the ICL area. Error bars represent mean ± SD. One-way ANOVA, multiple comparisons **p=0.0059, ****p<0.0001. n=9-12 AOB sections from 3-4 mice per treatment.

### ArcTRAP+ IGCs show increased intrinsic excitability for days after a single resident-intruder interaction

Previous studies indicated that Arc-positive IGCs display changes in their intrinsic excitability 4-8 hours after a single resident-intruder assay (Cansler, et al., 2017) or mating (Gao, et al., 2017). Here, we sought to understand how these changes in intrinsic excitability were affected over a one-week time-course. We hypothesized that if Arc-positive IGCs were playing a role in regulating behaviorally relevant incoming chemosensory information, we would see some persistent changes in excitability across a longer time period. Resident mice were TRAPed on Day 0 via a resident-intruder interaction with a BALB/cJ male. Following this single resident-intruder assay, acute slices were prepared for whole cell patch clamp electrophysiology on day 1, 3, 5, or 7. We recorded a series of electrophysiological challenges in current clamp (I-Clamp) and voltage clamp (V-Clamp) from tdTomato-positive (ArcTRAP+) and tdTomato-negative (ArcTRAP-) IGCs. Sample I-Clamp traces from ArcTRAP+ and ArcTRAP- IGCs on day 1, 3, 5, and 7 can be seen in Figure 2B. We measured 23 electrophysiological features from I-Clamp and V-Clamp challenges, which we have previously used to assist in identifying differing qualities within and across AOB interneuron types (Maksimova, et al., 2019). Previous results obtained during a 4-8 hour time window following social behavior indicated that maximal spiking frequency increased in Arc-expressing IGCs (Cansler, et al., 2017). We therefore investigated whether ArcTRAP+ and ArcTRAP- IGCs also demonstrated differences in spiking frequency (Fig. 2C-D). We found that the ArcTRAP+ population was more excitable than the ArcTRAP-population (one-way ANOVA, p=0.016, Fig. 2C-D). We investigated other electrophysiological features across fluorescence type and test day (Fig. 2E-F). We also found a difference in spontaneous excitatory post synaptic current (EPSC) amplitudes (two-way ANOVA, main effect of test day, p=0.047; Fig. 2E), a feature which was not different in the Arc-d4EGFP+ and Arc-d4EGFP- populations over the 4-8 hour timeline (Cansler et al., 2017). We did not see a difference in the hyperpolarization-activated (I_H_) currents, a notable feature of all IGCs (Hu, et al., 2016). These data indicated that differences in ArcTRAP+ IGCs persisted across a long, behaviorally relevant time scale. We next wanted to test whether reinforcing the behavior experience through repeated resident-intruder interactions would have an effect on the electrophysiological phenotypes of these cells.

**Figure 2.**
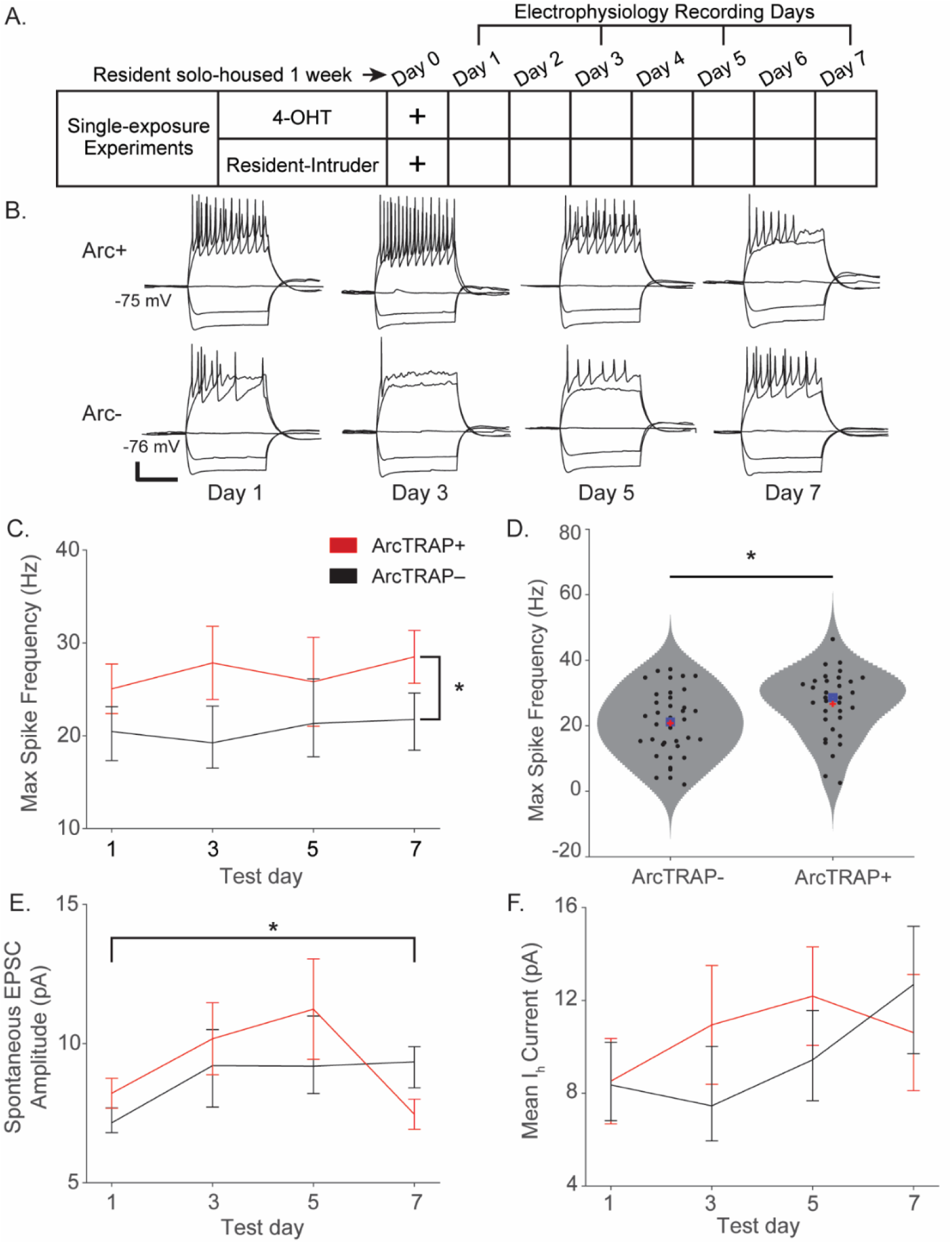
ArcTRAP+ IGCs show increased intrinsic excitability for 1-3 days following a single resident-intruder interaction. (**A**) Depiction of resident-intruder interaction followed by electrophysiology recordings on either day 1, 3, 5, or 7. (**B**) Sample current clamp traces from ArcTRAP+ and ArcTRAP- IGCs on days 1, 3, 5, and 7. Scale bar represents 0.25 seconds and 20mV. (**C**) Max spiking frequency of Arc+ and Arc- IGCs, split across days. Error bars: mean ± SEM. Two-way ANOVA, main effect of fluorescence p=0.018. (**D**) Max spiking frequency of ArcTRAP- and ArcTRAP+ IGCs, grouped from days 1, 3, 5, and 7. One-way ANOVA, p=0.016. Red “+” indicates mean, blue square indicates median. (**E**) Spontaneous EPSC amplitude of ArcTRAP+ and ArcTRAP- IGCs, split across days. Error bars: mean ± SEM. Two-way ANOVA, main test day effect, p=0.047. (**F**) Mean I_H_ current of ArcTRAP+ and ArcTRAP- IGCs, split across days. Error bars: mean ± SEM. Two-way ANOVA, ns.

### A similar population of IGCs re-express Arc upon re-exposure to the same intruder mouse

Given their days-long increases in excitability, we hypothesized that Arc-expressing AOB IGCs may contribute to experience-dependent chemosensory learning. For example, Arc-expressing IGCs may inhibit the population of AOB mitral cells activated during a salient social behavior, like aggression, and continue shaping mitral cell activation through repeated interactions. If Arc-expressing IGCs were indeed contributing to social behavior plasticity, we hypothesized that similar populations of IGCs would become activated during repeated behavior interactions with the same mouse. To test this hypothesis, we utilized triple-transgenic ArcTRAP-d4EGFP mice. In addition to the tamoxifen inducible Cre and tdTomato reporter (ArcTRAP), these mice harbor a bacterial artificial chromosome (BAC) that contains a transiently active EGFP induced by the Arc promotor (Arc-d4EGFP) (Grinevich, et al., 2009). These triple-transgenic mice allow us to permanently label cells expressing Arc on Day 0 with tdTomato, and then transiently label Arc-expressing cells on subsequent behavioral testing days with d4EGFP (Fig. 3A). Male resident ArcTRAP-d4EGFP were TRAPed on day 0 with a male BALB/cJ intruder. On days 1 and 2, the resident was re-exposed to the same BALB/cJ intruder in the resident-intruder assay. On day 3, resident mice were either exposed to the same BALB/cJ intruder seen on the previous days, or introduced to a female BALB/cJ. Three hours after the final resident-intruder assay on day 3, acute slices were prepared for 2-photon imaging. Colocalization analysis was performed to determine how much overlap existed between the tdTomato and d4EGFP signals in male-male and male-female exposed mice (Fig. 3B). Residents that were exposed to the same male mouse on all 4 days of the testing had increased colocalization of tdTomato and d4EGFP (Unpaired Student’s *t*-test, d4EGFP/tdTomato p=0.038, tdTomato/d4EGFP p<0.0001) compared to residents that were exposed to a male on the first three days and a female on the last day. These data show that IGCs re-express Arc after multiple social experiences with the same mouse, and that the chemosignals produced by male versus female mice activate largely non-overlapping populations of IGCs in the AOB.

**Figure 3.**
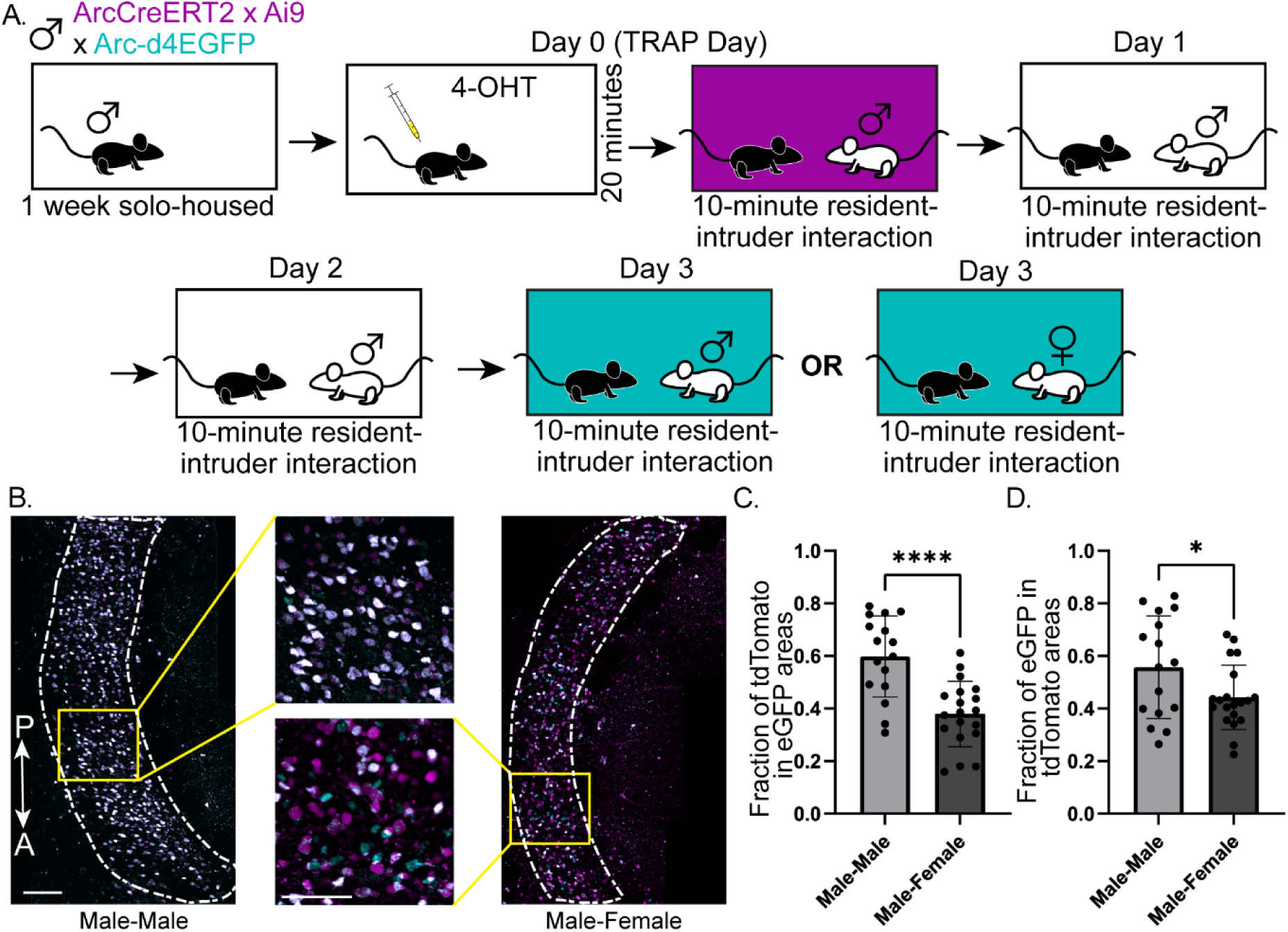
Re-exposure to the same intruder induces Arc expression in a similar population of IGCs across multiple days. (**A**) Depiction of repeated resident-intruder behavioral paradigm involving triple transgenic (ArcTRAP x Ai9 x Arc-d4EGFP) mice. 4-OHT injection was paired with resident-intruder exposure on Day 0. On Day 3, resident mice were either exposed to the same male intruder seen in the previous days, or to a novel female intruder. (**B**) Left: AOB image from a resident exposed to the same male intruder on all 4 days. Right: AOB image from a resident exposed to the same male intruder for three days, and a novel female intruder on the fourth day. Top middle: ICL inset from male-male exposure. Bottom middle: ICL inset from male-female exposure. tdTomato (magenta) label induced by Arc expression on Day 0. GFP (cyan) label induced by Arc expression on Day 3. White indicates colocalization between tdTomato and EGFP. Dashed line outlines the ICL. Scale bars 100 µm. (**C**) Colocalization analyses for the fraction of tdTomato+ pixels (magenta) colocalized with eGFP (cyan). (**D**) Colocalization analysis for the fraction of eGFP pixels colocalized with tdTomato pixels. Unpaired Student’s *t*-test, *p=0.038, ****p<0.0001. n=16-19 AOB slices from 4-5 animals.

### ArcTRAP+ IGCs sustain increased intrinsic excitability across daily resident-intruder interactions

Having shown that ArcTRAP+ IGC excitability is elevated for days following a single resident-intruder interaction, and that ArcTRAP+ IGCs are re-activated by repeated interactions with the same animal, we proceeded to investigate the physiological responses of ArcTRAP+ IGCs during repeated resident-intruder interactions. If ArcTRAP+ IGCs show increased excitability during repeated social encounters, this would further suggest that these cells are capable of supporting AOS-mediated social behavior plasticity. To test this, we conducted similar electrophysiology experiments to those described in Figure 2. However, in this cohort, we re-exposed resident mice to the same intruder every day leading up to the day of electrophysiology recording, including the day of recording (Fig. 4A). On day 3, 5, or 7, acute slices were prepared for whole cell patch clamp electrophysiology. The same I-Clamp and V-Clamp challenges were applied to ArcTRAP+ and ArcTRAP- IGCs (Fig. 4B). We found that ArcTRAP+ IGCs had a significantly increased maximum spiking frequency compared to ArcTRAP- IGCs (one-way ANOVA, p=0.013; Fig. 4C-D). We did not see a significant difference in spontaneous EPSC amplitudes when the mice underwent repeated resident-intruder exposures (Fig. 4E). Significant increases in I_H_ currents were found with re-exposure in the ArcTRAP+ population (two-way ANOVA, main effect of fluorescence, p=0.025; Fig. 4F), which were not seen with single exposure. These results indicate that ArcTRAP+ IGCs maintain increased intrinsic excitability features across multiple behavior exposures, potentially suggesting a role in AOS-mediated behavioral plasticity.

**Figure 4.**
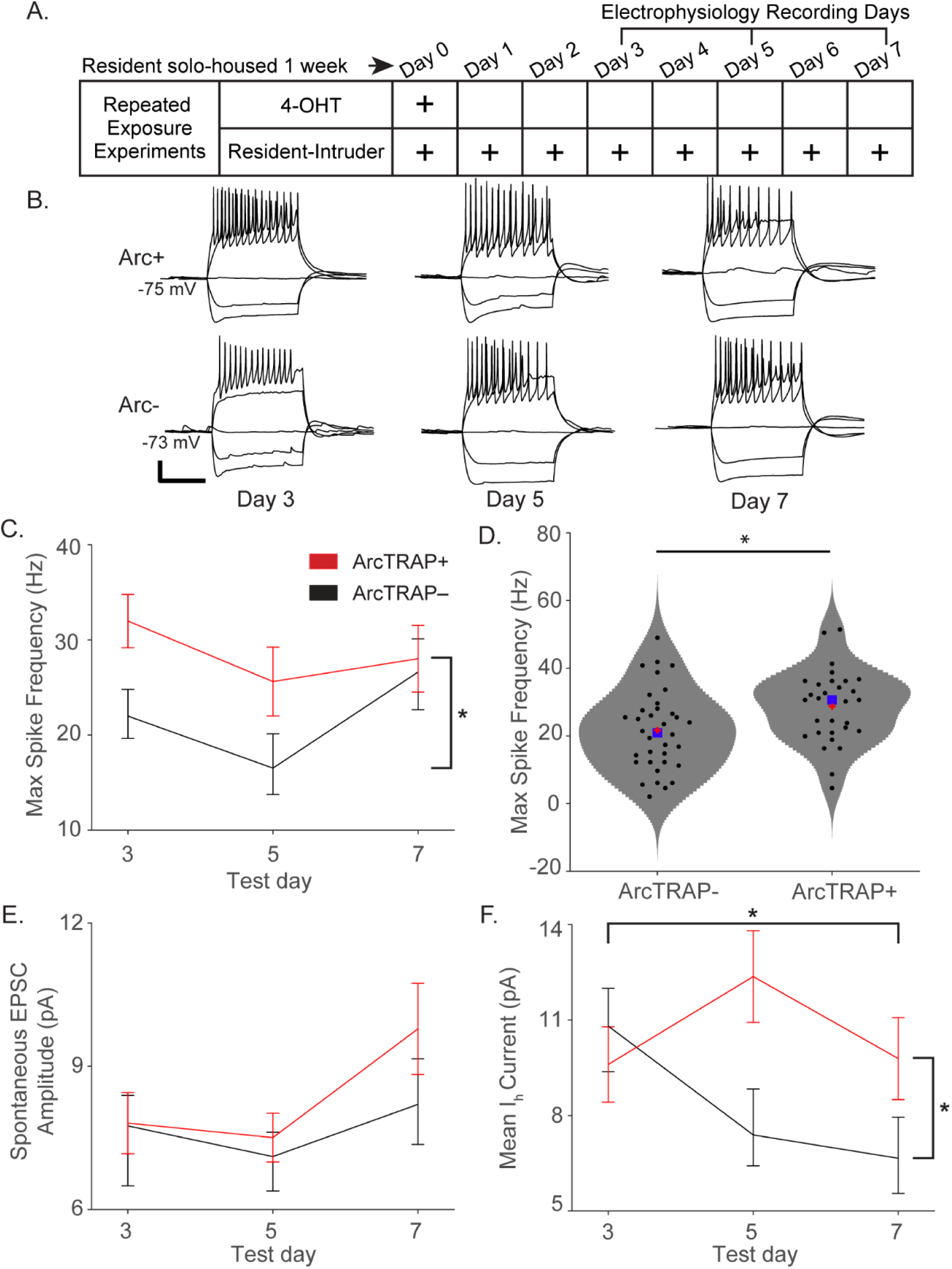
ArcTRAP+ IGCs show increased intrinsic excitability after repeated resident-intruder interactions. (**A**) Depiction of resident-intruder interactions followed by electrophysiology recordings on either day 3, 5, or 7. (**B**) Sample current clamp traces from ArcTRAP+ and ArcTRAP- IGCs on days 3, 5, and 7. Scale bar represents 0.25 seconds and 20mV. (**C**) Max spiking frequency of ArcTRAP+ and ArcTRAP- IGCs, split across days. Error bars: mean ± SEM. Two-way ANOVA, main fluorescence effect p=0.014. (**D**) Max spiking frequency of ArcTRAP- and ArcTRAP+ IGCs, grouped from days 3, 5, and 7. One-way ANOVA, p=0.013. Red “+” indicates mean, blue square indicates median. (**E**) Spontaneous EPSC amplitude of ArcTRAP+ and ArcTRAP- IGCs, split across days. Error bars: mean ± SEM. Two-way ANOVA, ns. (**F**) Mean I_H_ current of ArcTRAP+ and ArcTRAP- IGCs, split across days. Error bars: mean ± SEM. Two-way ANOVA, main fluorescence effect p=0.025. Combined test day x fluorescence effect p=0.047.

The changes in ArcTRAP+ IGC physiology we saw in both single and repeated resident-intruder interactions were largely consistent with those observed in d4EGFP-expressing cells 4-8 hours after resident-intruder interactions (Cansler, et al., 2017). Previous work indicated that IGCs are electrophysiologically heterogenous (Cansler, et al., 2017; Gao, et al., 2017; Maksimova, et al., 2019) but that they can be readily distinguished from other AOB IGCs based on multidimensional analysis of electrophysiological parameters (Maksimova, et al., 2019). In order to more systematically investigate the impacts of these behavioral conditions on ArcTRAP+ and ArcTRAP- IGCs, we analyzed 137 IGCs recorded in both “Single exposure” and repeated exposure (“Re-exposure”) conditions (Fig. 5). Cluster analysis identified both a “high excitability” and a “low excitability” IGC population across all recordings (Fig. 5B). The low excitability cluster (Cluster 1) was enriched for ArcTRAP- IGCs across days and experiments (17/22 cells, p<0.01, binomial test), whereas the high excitability cluster (Cluster 3) was enriched for ArcTRAP+ IGCs (16/21 cells, p<0.01, binomial test). The vast majority of IGCs were not included in either of these clusters (94/137, p=0.54, binomial test), suggesting experience-dependent excitability changes may ride on top of a highly heterogenous baseline. In order to determine whether ArcTRAP+ cells are shifted towards the high excitability state, we used linear discriminant analysis (LDA) to generate a multidimensional vector (Eigenvector) that best distinguished high-excitability and low-excitability clusters (Fig. 5C). Projecting each recording along this primary LDA Eigenvector allowed us to quantify the relative closeness of each IGC to the high-excitability or low-excitability state (Fig. 5D-E). This comparison revealed strong separation of ArcTRAP+ and ArcTRAP- IGCs in both “Single exposure” (SE) and “Re-exposure” (RE) conditions (Fig. 5D). Furthermore, this comparison confirmed that ArcTRAP+ IGC excitability increases were present at 1 and 3 days post-TRAPing in SE experiments, and that these increases were present at 3 and 5 days post-TRAPing in RE experiments (Fig. 5E). In day 7 RE experiments, we observed no difference in ArcTRAP+ and ArcTRAP- excitability (p=0.71, Kruskal-Wallis test). This appeared to be due to an increase in ArcTRAP-excitability, rather than a decrease in ArcTRAP+ excitability (Fig. 5C). Given that each resident-intruder interaction activates some IGCs that were not activated on “TRAP Day” (Fig. 3), this lack of separation on day 7 may reflect the inclusion of cells that began expressing Arc on days 1-7 (post TRAP), and therefore did not express tdTomato at the time of recording. Overall, these results confirm and extend the observations that Arc-expressing AOB IGCs shift toward high excitability states for days following resident-intruder interactions, in both single exposure and re-exposure conditions.

**Figure 5.**
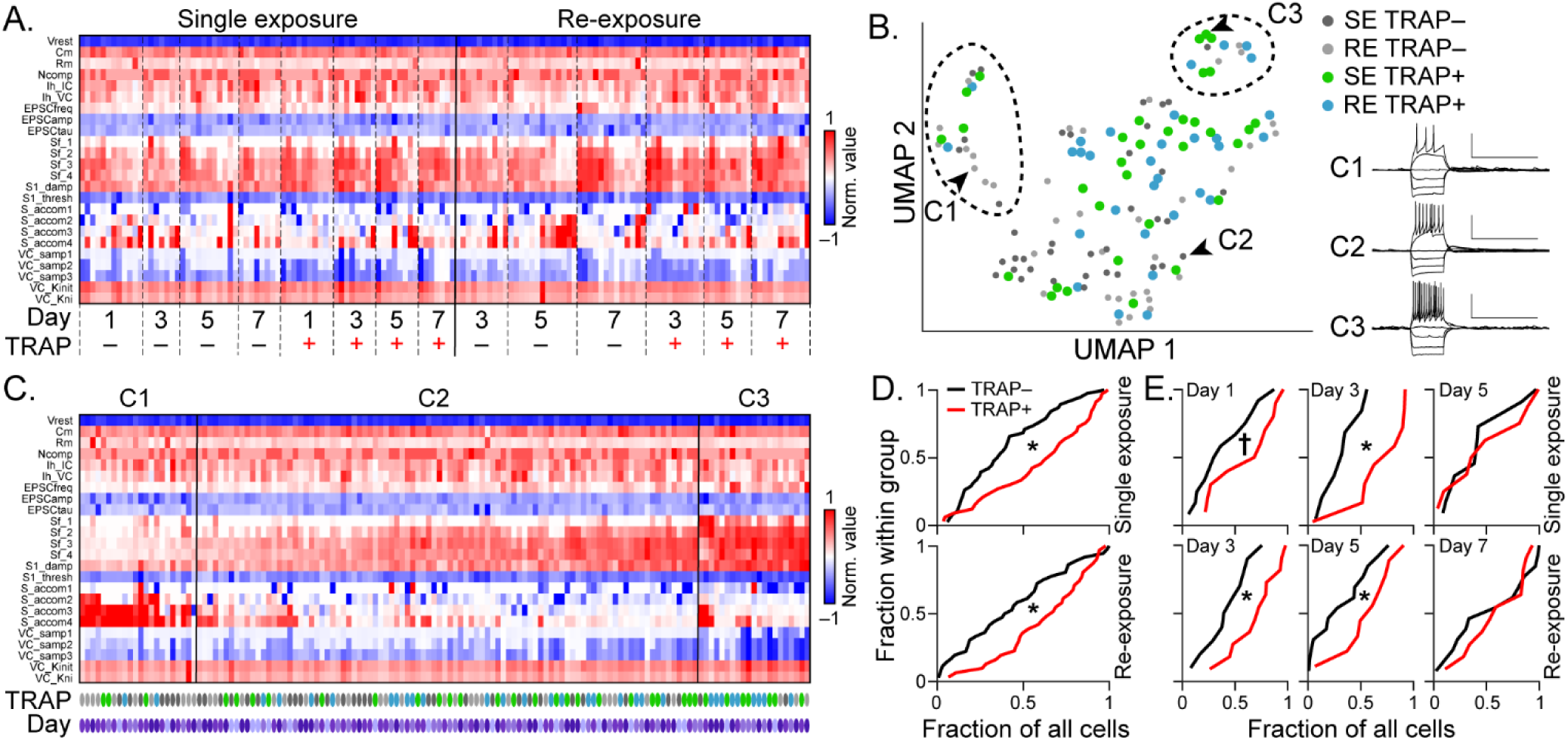
Multidimensional analysis of IGC electrophysiological properties. (**A**) Heatmap of 23 electrophysiological properties recorded for 137 IGCs included in Single exposure (SE) and Re-exposure (RE) experiments (Figs. 2 and 4, respectively). Measurements were normalized by the maximum value observed and colorized based on the sign of the measurement for display purposes. (**B**) Uniform manifold approximation and projection (UMAP) of all cells in Panel A, colorized by their identity. Despite high levels of heterogeneity, cluster analysis identified IGCs with low excitability (C1), high excitability (C3), and intermediate excitability (C2). Insets show cells identified by arrowheads. Scale bars: 50 mV, 1 s. (**C**) Clustered heat map with cells displayed based on their projection along an Eigenvector that best separates cells in Cluster 1 (low excitability) from cells in Cluster 3 (high excitability). Day of measurement is indicated by purple ovals (Day 1: lightest, Day 7: darkest) and TRAP condition is indicated using the same color codes as in Panel B. (**D**) Cumulative distributions of TRAP+ (red) and TRAP– (black) IGCs along the Eigenvector indicated in Panel C for Single exposure (top) and Re-exposure (bottom) experiments. (**E**) Relative ranks of TRAP+ and TRAP– IGCs along LDA Eigenvector, broken down by experimental day. * indicates p < 0.05, † indicate p < 0.1, Kruskal-Wallis test.

### Aggressive behavior in male mice increases across repeated resident-intruder interactions

When conducting the repeated exposures to the resident-intruder assay, we noted changes in the resident’s aggression levels over time. Given that intermale aggression is AOS-mediated (Stowers, et al., 2002), we investigated whether this aggression is supported by experience-dependent plasticity in the AOB. Other studies have indicated that male resident mice tend to become more aggressive when undergoing short duration, repeated resident-intruder interactions (Sunkin, et al., 2013). Because of this, we wanted to identify whether male ArcTRAP resident mice increase their aggressive behavior across repeated, 10-minute resident-intruder interactions once per day for eight days. Video was recorded during the repeated resident-intruder interactions for electrophysiology. We manually quantified the behavior by counting the number of resident-initiated aggressive interactions, as well as the latency to the first aggressive interaction (Fig. 6B-C). We found that resident mice significantly increased the number of aggressive interactions toward the end of the eight-day behavior period (days 5, 6 and 7) (one-way ANOVA, p<0.05). In line with an increase in aggression, we also saw a decrease in the latency to the first attack, with a significant decrease in latency between Day 3 and Day 5 of testing (one-way ANOVA, p=0.0075). These results indicate that experimental conditions that re-activate ArcTRAP+ AOB IGCs coincide with changes in an AOS-mediated social behavior.

**Figure 6.**
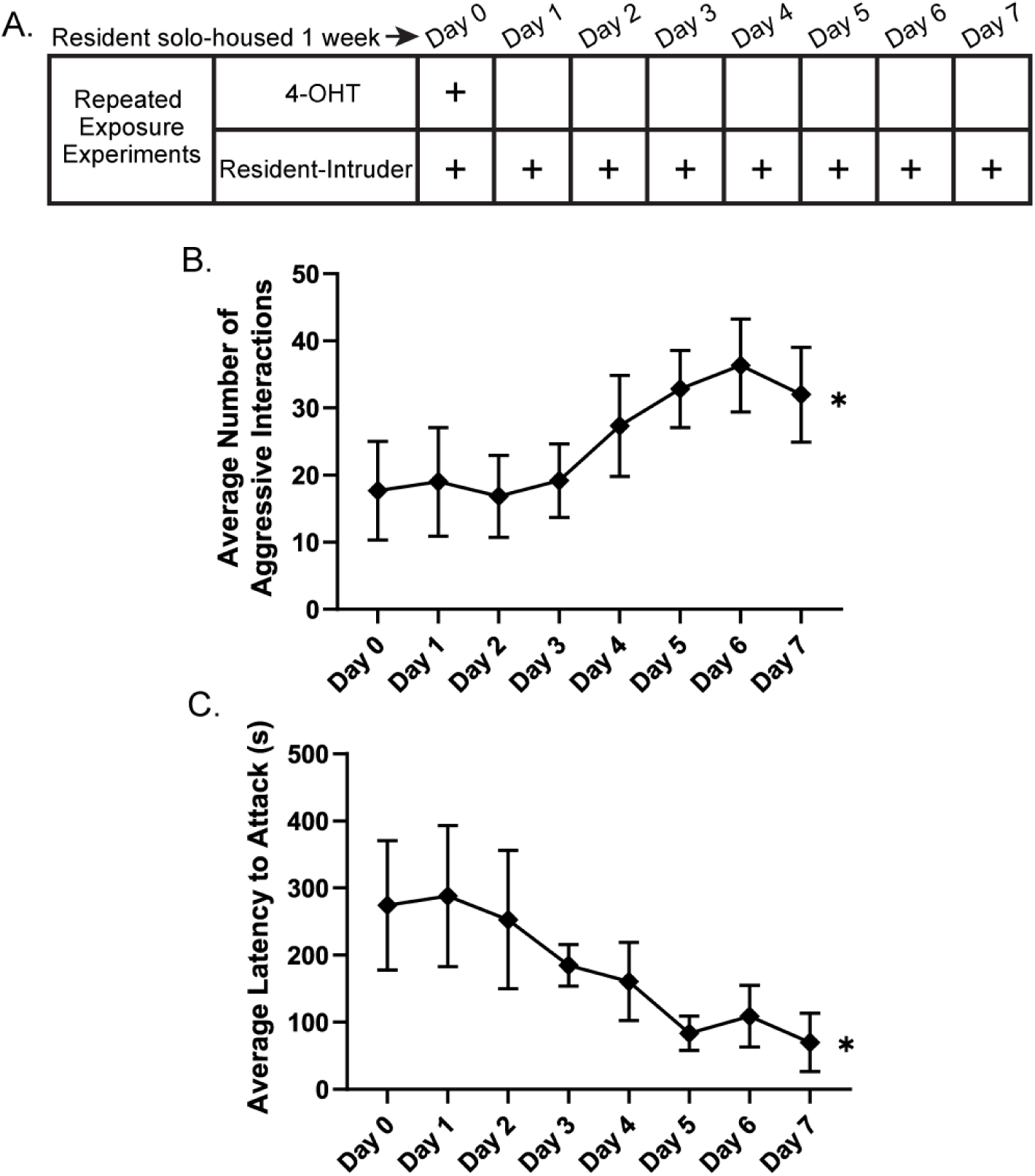
Ramping aggression in ArcTRAP males during repeated resident-intruder interactions. (**A**) Depiction of resident-intruder schedule across eight days. (**B**) Average number of resident-initiated aggressive interactions across repeated resident-intruder interactions. (**C**) Average latency to first attack across repeated resident-intruder interactions. Error bars represent mean ± SEM. One-way ANOVA. *p<0.05, **p=0.0075, n=6.

### Inhibiting ArcTRAP+ IGCs with DREADDs abolishes the intermale ramping aggression

We next sought to test whether ArcTRAP+ IGCs participate in intermale aggressive ramping. We hypothesized that inhibiting the ArcTRAP+ population may affect these behavioral outputs of chemosensory information processing. To test this hypothesis, we utilized inhibitory designer receptors exclusively activated by designer drugs (G_i_-DREADDs). A Cre-dependent AAV containing the G_i_-DREADD was injected into the AOB of male resident ArcTRAP mice. Control mice received a Cre-dependent AAV-mCherry. After one week of recovery from surgery, while also being solo-housed, the mice were TRAPed on Day 0, activating Cre and inducing expression of the G_i_-DREADD (or mCherry-only control). Following this initial resident-intruder interaction, residents were solo-housed for an additional week to allow the G_i_-DREADD receptor to be strongly expressed. Residents then underwent repeated resident-intruder interactions once per day for seven days and received either a clozapine-n-oxide (CNO) or dimethylsulfoxide (DMSO; vehicle) injection 20 minutes prior to behavior (Fig. 7A). To ensure that the G_i_-DREADD was having the anticipated hyperpolarizing effect on IGCs, we conducted a drug wash-on electrophysiology experiment in AOB slices (Fig. 7B). Resting membrane potential was recorded for a 32-minute duration. Compared to IGCs expressing only mCherry, the G_i_-DREADD expressing IGCs showed a significant hyperpolarization induced by the CNO wash-on, which persisted through the post-drug period (two-way ANOVA, p<0.01), confirming that the G_i_-DREADDs were having the anticipated hyperpolarizing effect in IGCs.

**Figure 7.**
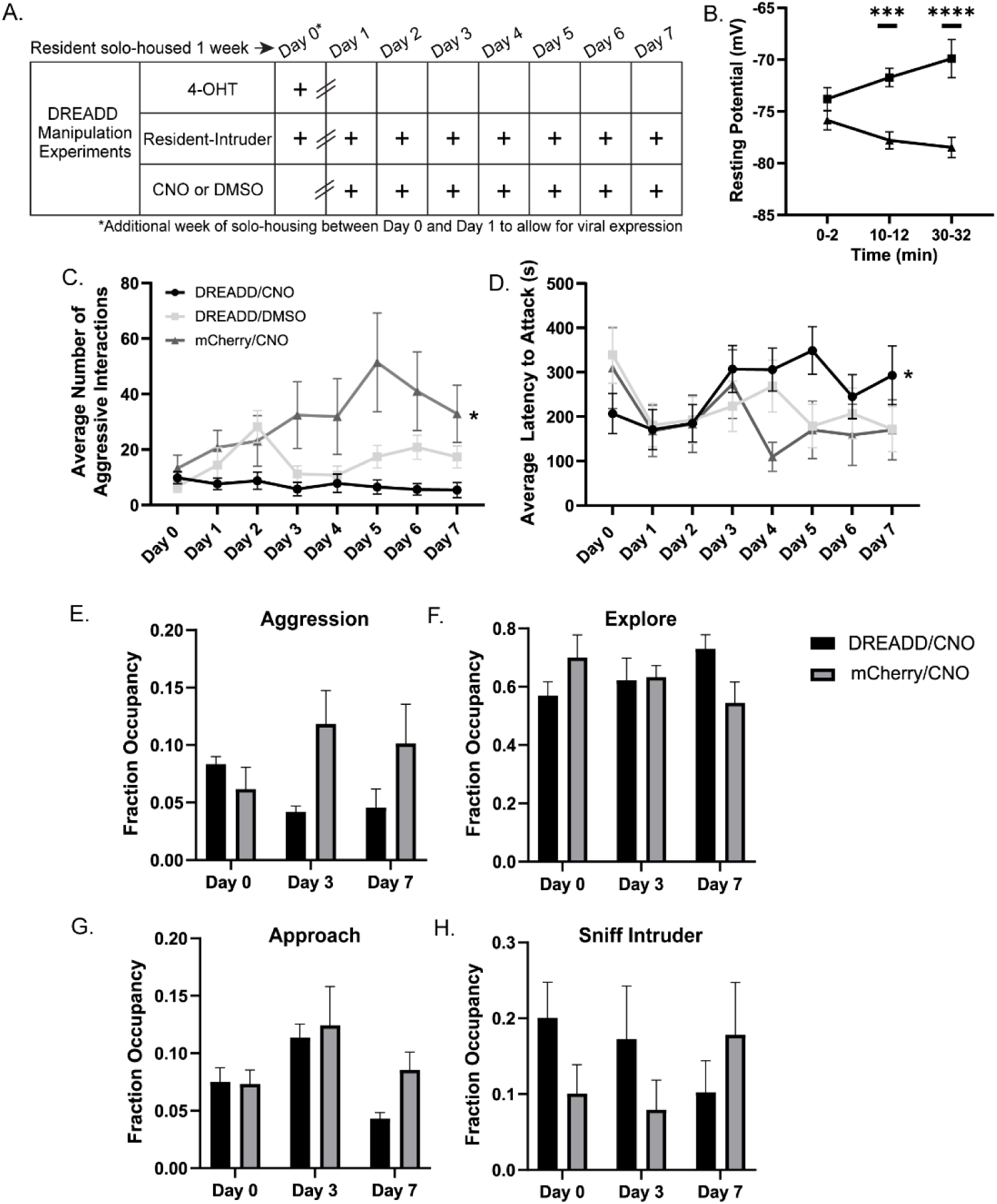
Inhibiting ArcTRAP+ IGCs with a Gi-DREADD abolishes ramping aggression in male mice. (**A**) Depiction of the resident-intruder interaction schedule with drug injections. Mice were solo-housed for one week prior to Day 0, and for an additional week after Day 0. Subsequently, they received daily CNO or DMSO (control) injections before the resident-intruder paradigm for the duration of the experiment. (**B**) Whole-cell patch clamp membrane potential recordings from IGCs in AOB slices during CNO wash-on. IGCs expressing either an AAV-Gi-DREADD or AAV-mCherry were patched and recorded from for a period of 32 minutes. Baseline resting potential was measured from minutes 0-2. A 10-minute CNO wash-on period occurred between minutes 2-12. A 20-minute wash-off period occurred between minutes 12-32 (though no wash-off effect was seen). Error bars: mean ± SD. Two-way ANOVA, multiple comparisons, **p<0.01. n=6-17 cells. (**C**) Average number of aggressive interactions per treatment group across eight days. Two-way ANOVA, multiple comparisons. *p<0.02 between DREADD/CNO and mCherry/CNO groups. n=9-13 mice. (**D**) Average latency to attack per treatment group across eight days. Two-way ANOVA, multiple comparisons. *p=0.0082 between DREADD/CNO and mCherry/CNO. n=9-13 mice. (**E-F**) DeepLabCut occupancy analysis for listed behaviors. Subset of animals used in panel C, D. n=4-6 mice. Error bars represent mean ± SEM.

After verifying that the G_i_-DREADD was actively inducing hyperpolarization in ArcTRAP+ IGCs during a CNO application, we assessed the effect of hyperpolarizing ArcTRAP+ IGCs on behavioral outcomes. We manually measured the number of resident-initiated aggressive interactions and latency to first attack, finding that mice receiving the AAV-mCherry and CNO injection displayed the same aggressive ramping behavior (Fig. 7C) as seen in unmanipulated residents (Fig. 6B). They also showed a similar decrease in latency to attack (Fig. 7D). Consistent with our hypothesis that ArcTRAP+ IGCs participate in ramping intermale aggression, we found that mice receiving the G_i_-DREADD and CNO injection lacked the ramping aggression seen in the mCherry control mice (Fig. 7C). They also displayed an increased latency to first attack (Fig. 7D). Mice receiving the G_i_-DREADD and DMSO control injection displayed moderately increased aggression, and increased latency to first attack, however they were not at the same levels of the mCherry control (Fig. 7C-D). This may be explained through physiological changes induced solely by the expression of a G_i_-DREADD, in the absence of the activating drug, which has been shown in other studies (Saloman, et al., 2016). Altogether, these data suggest that proper regulation of signaling in inhibitory IGCs is necessary for sex-typical intermale ramping aggression in the resident-intruder paradigm.

To determine whether these manipulations altered other aspects of social behavior, on a subset of experiments (for which we had top-down video recordings), we used the machine learning-based DeepLabCut software (Mathis, et al., 2018) and random forest-based behavioral segmentation. This analysis showed that control mCherry/CNO mice had increased aggression on Days 3 and 7 compared to DREADD/CNO mice (Fig. 7E). We did not see a difference in other relevant behaviors, such as exploration (time spent exploring the cage, sniffing bedding, etc.), approach (resident approaching intruder), or sniffing behavior (resident sniffing intruder) (Fig. 7E-H). We did observe a trend in the data suggesting that the DREADD/CNO group demonstrated higher non-aggressive sniffing behaviors than controls (Fig. 7H). Combined, these data show that although expression and activation of G_i_-DREADDs strongly decreases the levels of aggression in resident mice, other non-aggressive behaviors are largely unchanged. This supports our hypothesis that ArcTRAP+ IGCs play a role in specifically regulating AOS-mediated aggressive behavior in the context of the resident-intruder paradigm.

## Discussion

Transgenic silencing and physical ablation of the VNO established that the AOS is essential for several aspects of rodent social behavior (Keller, et al., 2006; Stowers, et al., 2002). The AOB itself has also been targeted for selective manipulation in the context of social behavior (Brennan, et al., 1990; Kunkhyen, et al., 2017; Otsuka, et al., 2001). Despite the increased availability of cell type-specific transgenic tools for neuroscience, there are few tools that are selective enough to help delineate the functions of the several classes of neurons in the AOB (Cansler, et al., 2017; Gao, et al., 2017; Maksimova, et al., 2019; Zhang, et al., 2020). The availability of transgenic tools for labeling and manipulating Arc-expressing neurons is especially helpful for studying AOB circuit function, as IGCs are the predominant cell type expressing this immediate-early gene in the AOB (Cansler, et al., 2017; Gao, et al., 2017; Matsuoka, et al., 2002). Building on previous work that showed Arc-expressing IGCs demonstrate experience-dependent plasticity, we sought to understand if this plasticity participated in longer-term AOS function, and whether the activity of these cells ultimately affects AOB signaling and the subsequent behavioral output.

ArcTRAP mice (Guenthner, et al., 2013) provided an opportunity to investigate these questions. However, these and other tools that enable transient IEG-based labeling have limitations, requiring validation in a circuit and cell type-specific manner (Allen, et al., 2017; Guenthner, et al., 2013; Reijmers, et al., 2009). We investigated the utility of the ArcTRAP tool for studying AOB IGCs in the context of intermale territorial aggression, a context that has already been shown to induce robust Arc expression in AOB IGCs (Cansler, et al., 2017). This behavioral context is certainly not the only condition in which IGCs express Arc (Gao, et al., 2017; Matsuoka, et al., 2002), but it is a well-established AOS-mediated behavior, making it an attractive platform in which to perform initial testing and validation. TRAPing in this context was robust and cell type-specific (Fig. 1). Although TRAPing was strongest in social chemosensory conditions, we also observed increased tdTomato labeling in mice only receiving 4-OHT in their home cage (Fig. 1). In other brain regions, IEGs like Arc are not simply induced by neuronal excitation but also by novelty (Cleland, et al., 2017). Our results may indicate that modest sampling of self chemosignals is sufficient to drive Arc-CreERT2-mediated recombination, or perhaps that AOB IGCs have spontaneous activity levels capable of driving Arc expression. Regardless, these initial studies confirmed that ArcTRAP-based approaches selectively label AOB IGCs, and that the labeling reflects the chemosensory conditions present in the time window following 4-OHT injection as expected.

Two recent studies showed that Arc-expressing IGCs have increased excitability compared to other IGCs, and a major component of this effect was a change in the ability for Arc-expressing IGCs to maintain high spiking frequencies in the presence of strong depolarization (Cansler, et al., 2017; Gao, et al., 2017). We found that ArcTRAP+ IGCs maintain similar increases in excitability for 3 days following a single resident-intruder interaction (Fig. 2). As in previous studies, the increased intrinsic excitability of ArcTRAP+ IGCs was not accompanied by alterations in spontaneous EPSC frequency or amplitude (Fig. 2E). The lack of obvious synaptic plasticity in these conditions does not preclude a subtle change in synaptic properties, but these experiments were not designed to tease apart such nuanced differences. Future studies would be needed to identify, for example, whether ArcTRAP+ IGCs are primed for future plasticity (“metaplasticity”) as has been seen in other Arc-expressing cells (Jakkamsetti, et al., 2013). The observation that ArcTRAP+ IGCs maintain elevated excitability for several days after a social chemosensory encounter indicated that these cells are modified over time courses compatible with storing a chemosensory memory, a topic of much interest over the past several decades (reviewed in Brennan, 2009). To determine whether Arc-expressing IGCs might play roles in chemosensory memory and/or social behavior plasticity, we proceeded to investigate Arc expression patterns after multiple social encounters.

In the hippocampus, well-known for its role in spatial and episodic memory, re-exposure to a familiar environment elicits Arc expression in a similar population of cells (Chawla, et al., 2005). In contrast, exposure to a novel environment induces Arc expression in a mostly non-overlapping neuronal population (Chawla, et al., 2005). If Arc-expressing AOB IGC ensembles possess similar qualities in social chemosensory contexts, this could suggest the presence of an inhibitory “memory trace” at the first stage of sensory processing. Re-exposing a resident male to a familiar male intruder for 3 days after TRAPing induced highly overlapping Arc expression among ArcTRAP+ IGCs (Fig. 3). However, when a novel female mouse was introduced on the fourth day, Arc was expressed in a mostly non-overlapping population of IGCs (Fig. 3). These data indicated that a similar ensemble of IGCs are re-activated, re-express Arc, across multiple social interactions, and that the population of IGCs that express Arc reflect the social chemosensory environment. This also indicates that cells expressing Arc do not chronically express Arc, but that they transiently express Arc as a result of the specific intruder-induced AOB excitation.

It is not clear whether the re-expression of Arc across multiple days would sustain existing elevations of intrinsic excitability, or whether this would produce additional physiological effects. We therefore tested whether Arc re-expression produced new alterations to IGC physiology. If physiological changes in ArcTRAP+ IGCs scaled with each experience, we would expect to see an enhancement of physiological features that were affected after additional chemosensory encounters. We did find that increased IGC excitability persisted in ArcTRAP+ and ArcTRAP- IGCs following multiple resident-intruder encounters. However, this increased excitability in “re-exposure” experiments did not scale with repeated social encounters (Fig. 4). In both the single-exposure and re-exposure paradigms, we found that maximum spiking frequency of ArcTRAP+ and ArcTRAP- groups was statistically indistinguishable on Day 7. However, in the single-exposure paradigm, ArcTRAP+ cells reduced their excitability to control levels by Day 7, suggesting that, in the absence of reactivation, the excitability effect waned. In contrast, in the re-exposure paradigm, we found that the similarities between ArcTRAP+ and ArcTRAP- maximum spiking frequency on Day 7 were driven by an increase in excitability in the ArcTRAP- group. We hypothesize that this may have been due to ArcTRAP- IGCs (cells that were not TRAPed on Day 0) being stimulated to express Arc between Days 1-7. These cells would not express the tdTomato marker (because Arc expression occurred outside the 4-OHT time window), but would have expressed Arc in subsequent days, and therefore increased their excitability. This hypothesis is consistent with observations made during repeated exposure experiments involving ArcTRAP-d4eGFP mice (Fig. 3). In these experiments, we saw that a majority of ArcTRAP+ IGCs re-expressed Arc, but some ArcTRAP+ IGCs did not re-express Arc. Other IGCs expressed Arc on the last day, but not the first (Fig. 3). Future electrophysiology studies will be needed to determine whether more nuanced changes in IGC physiology, morphology, or gene expression occur as IGCs repeatedly express Arc across multiple social encounters.

IGCs are a heterogeneous population electrophysiologically, morphologically, and genetically (Cansler, et al., 2017; Larriva-Sahd, 2008; Maksimova, et al., 2019). This heterogeneity may have several sources, including cellular maturity; IGCs are continuously added to the AOB via adult neurogenesis in the subventricular zone and migration via the rostral migratory stream (Bonfanti, et al., 1997). We explored the electrophysiological diversity of IGCs in the context of both single-exposure and re-exposure experiments in order to place ArcTRAP+ and ArcTRAP- excitability changes in a broader context (Fig. 5). Clustering analysis identified high and low excitability groups that were enriched in ArcTRAP+ and ArcTRAP- IGCs, respectively. Using these two clusters as templates, we used a single multidimensional vector that best separated these two populations. We used this Eigenvector as a proxy for the complement of physiological changes that AOB IGCs demonstrate in these conditions, which may relate to their computational functions within the circuit (Fig. 5). This analysis showed that ArcTRAP+ and ArcTRAP- IGCs do not acquire distinct electrophysiological phenotypes. Rather, ArcTRAP+ IGCs are seemingly pushed toward a high excitability phenotype. The circuit functions of lower- and higher-excitability IGCs are not yet known, and future studies will be needed to determine how Arc expression impacts cellular maturity and longevity.

In observing the interactions between males during re-exposure electrophysiology experiments, we noted that resident mice became more aggressive over the course of 7 days of re-exposure (Fig. 6). Given that ArcTRAP+ IGCs have increased excitability across this same time-course, we hypothesized that IGC activity may be playing a role in regulating these behavioral changes. When we inhibited the AOB ArcTRAP+ population with G_i_-DREADDs, we ablated the sex-typical aggressive ramping seen in repeated resident-intruder assays (Fig. 7). To our knowledge, this is the first report of an AOS-mediated behavioral effect driven by selective manipulation of an AOB interneuron subtype. The lack of ramping aggression in mice receiving G_i_-DREADDs and the DREADD agonist CNO could have been due to an off-target effect, for example a reduction in main olfactory function, or other behavioral deficits. However, a DeepLabCut-based investigation of the microstructure of these encounters showed that other socially relevant behaviors (exploring, sniffing, approaching), were not affected by the manipulation of ArcTRAP+ IGCs (Fig. 7). In animals expressing the G_i_-DREADD but receiving vehicle injections, we observed an intermediate phenotype between controls and CNO-injected animals (Fig. 7). This may indicate that expression of G_i_-DREADDs causes disruption of these cells in the absence of the selective ligand. Agonist-independent effects of G_i_-DREADD expression on neurons have been reported elsewhere, and represent an important consideration for these tools (Saloman, et al., 2016). Despite this possibility, the direction of the change is consistent with the overall conclusion of this experiment, which is that alteration of ArcTRAP+ IGC function reduces ramping aggression.

In these studies, we chose to selectively inhibit ArcTRAP+ IGCs, a manipulation likely to disinhibit AOB excitatory mitral cells. In female mice, AOB disinhibition causes failure of pregnancy maintenance, similar to the effects of a stranger male or its urine (Kaba, et al., 1988). If mitral cell hyperactivation increased the excitation by male chemosignals in the resident-intruder paradigm, one hypothesis would be that this would result in increased intermale aggressive behavior. However, the behavioral results indicate that this manipulation eliminated ramping aggression, an effect similar in nature to that of removing or silencing the VNO itself (Stowers, et al., 2002; Wekesa, et al., 1994). This suggests that Arc-expressing IGCs are necessary for the AOB to reliably transmit chemosensory information across repeated social encounters. In future studies, it will be important to investigate whether the manipulation of Arc-expressing IGCs similarly influence other AOS-mediated behaviors, such as mating and predator avoidance (Takahashi, et al., 2018; Takahashi, 2014).

In summary, these studies demonstrate that a population of plastic interneurons in an early chemosensory circuit display physiological features consistent with simple memory formation. These neurons, when silenced, result in disruption of natural escalation of aggressive behavior between males, a well-known chemosensory mediated behavior. Plasticity between excitatory and inhibitory interneurons is increasingly appreciated as critical for brain function, and these studies highlight that one of these roles is to support social behavior plasticity.

